# Genome engineering of *Nannochloropsis* with large deletions for constructing microalgal minigenomes

**DOI:** 10.1101/2020.10.08.332478

**Authors:** Qintao Wang, Yanhai Gong, Yuehui He, Yi Xin, Nana Lv, Xuefeng Du, Yun Li, Byeong-ryool Jeong, Jian Xu

**Affiliations:** Single-Cell Center, CAS Key Laboratory of Biofuels, Shandong Key Laboratory of Energy Genetics and Shandong Institute of Energy Research, Qingdao Institute of BioEnergy and Bioprocess Technology, Chinese Academy of Sciences, Qingdao, Shandong 266101, China; Qingdao National Laboratory of Marine Science and Technology, Qingdao, Shandong 266237, China; School of Energy and Chemical Engineering, Ulsan National Institute of Science and Technology, Ulsan 44919, Korea; University of Chinese Academy of Sciences, Beijing 100049, China

**Keywords:** oleaginous microalgae, *Nannochloropsis* spp., CRISPR-Cas system, genome editing, large genome fragment deletion

## Abstract

Industrial microalgae are promising photosynthetic cell factories, yet tools for targeted genome engineering are limited. Here for the model industrial oleaginous microalga *Nannochloropsis oceanica* we established a method to precisely and serially delete large genome fragments of ~100 kb from its 30.01-Mb nuclear genome. We started by identifying the “non-essential” chromosomal regions (i.e., low-expression region or LER) based on minimal gene expression under N-replete and N-depleted conditions. The largest such LER (LER1) is ~98 kb in size, located near the telomere of the 502.09 kb-long Chromosome 30 (Chr 30). We deleted 81 kb and further distal and proximal deletions of up to 110 kb (21.9% of Chr 30) in LER1 by dual targeting the boundaries with the episome-based CRISPR/Cas9 system. The telomere-deletion mutants showed normal telomeres consisting of CCCTAA repeats, revealing telomere regeneration capability after losing distal part of Chr 30. Interestingly, the deletions caused no significant alteration in growth, lipid production or photosynthesis (transcript-abundance change for < 3% genes under N depletion). We also performed double-deletion of both LER1 and LER2 (from Chr 9) that totals ~214 kb, and phenotypes are essentially normal. Therefore, loss of the large yet “non-essential” regions does not necessarily sacrifice important traits. Such serial targeted deletions of large genomic regions have not been reported in plants or microalgae, and will accelerate crafting minimal genomes as chassis for photosynthetic production.

## Introduction

Microalgae are photoautotrophic eukaryotic organisms that play a major role in the biogeochemical carbon cycling of our biosphere by assimilation of atmospheric CO_2_ (1, 2). In addition, microalgae have tremendous potential for producing biofuels, biomaterials and other platform chemicals in a renewable and sustainable manner while reducing greenhouse gas emission (3). However, realization of the potential requires extensive engineering of metabolism at the genetic and the genomic levels to maximize yields and minimize production costs (4, 5).

In general, genome is composed of many seemingly non-essential regions, which can be removed to create a “minimal genome”. For example, in higher eukaryotes, “junk" regions and/or unknown loci including transposons and repetitive elements can take up to 70% of the genome (6). Even in the compact bacterial genomes, the minimal genomes can be reduced to ~50-70% of the original size, based on the number of essential genes for normal growth if nutrients and stresses are not limiting (7, 8). Such “minimal genomes" can be employed as a chassis for building production strains, e.g., by introducing non-native biosynthetic pathways for target compounds (9). Notably, although a “minimal” genome of *Mycoplasma mycoides* has been synthesized by the bottom-up approach (10), *de novo* synthesis of eukaryotic genomes remain formidable due to their larger genome size and complexity (11). Therefore, top-down strategies that rationally determine and then delete non-essential regions from the native chromosome are attractive approaches for creating a minimal eukaryotic genome (12).

Deletion of target genomic regions can be achieved by various techniques, including the λ-red recombination system (13), Cre/loxP system (14), Flp/FRT system (15), Latour system (16), PCR-mediated chromosome splitting (17–19), gene replacement with meganuclease (20) and replacement-type recombination (21). In microalgae, however, such targeted deletions have hardly been successful due to their intrinsic problems. *Firstly*, genome-wide understanding that underlies rational selection and meaningful deletion of the target sites has been limited. *Secondly*, the efficiency of recombination is generally very low for microalgae (4, 22), despite a few examples of homologous recombination in microalgae (23–25), resulting in the lack of the aforementioned genome-deletion techniques.

Development of nuclease-based techniques is opening new possibilities for genetic manipulation of microalgae (4, 26). In particular, the clustered regularly interspaced short palindromic repeat (CRISPR)/Cas9 has been successfully employed in the microalgae such as *Chlamydomonas* (27–30), *Nannochloropsis* (26, 31–34), *Volvox carteri* (35), diatom (36–38), *Coccomyxa* sp (39) and *Euglena gracilis* (40), for gene knockout, knock-in, multiple knockout, homology-based small-fragment deletion of about 220 bp in *E. gracilis* (40). However, targeted deletions of large genomic fragments or regions have not been reported in microalgae (or plants), likely due to the generally low transformation efficiency of microalgae and the potentially harmful or even lethal effects of such deletions.

*Nannochloropsis* spp. are a phylogenetically distinct group of unicellular photosynthetic heterokonts that are widely distributed in sea, fresh and brackish waters. As Eustigmatophyceae they are more closely related to diatoms than to green algae. These heterokont microalgae are of industrial interest due to their ability to grow under a wide-range of conditions, and produce large amounts of lipids and high-value polyunsaturated fatty acids (PUFAs), e.g., eicosapentaenoic acid (41). Moreover, they are excellent research models for microalgal systems and synthetic biology, due to their small genome size and simple gene structure (42–47), as well as recently demonstrated genetic tools for *Nannochloropsis* (48), including overexpression (48–57), RNAi (58–60), multigene expression enabled by bidirectional promoters and ribosome skipping 2A sequences (24, 48, 61–63), markerless trait stacking through combined genome editing and marker recycling (34) and gene targeting via homologous recombination (24, 63–65).

*Nannochloropsis* spp. also feature extensive omics resources for functional assessment of chromosomal regions genome-wide. For example, in *Nannochloropsis oceanica* IMET1 which is an industrial strain for both TAG and EPA, rich resources of genomic (42, 47), transcriptomic (42, 66–69), proteomic (68–72), lipidomic (66, 67) as well as physiological data (73–77). Taking advantage of these resources, we employed this strain to establish a method to precisely and serially delete large genome fragments of ~100 kb from its 30.01-Mb nuclear genome. We started by identifying the “non-essential” chromosomal regions based on minimal gene expression under N-replete and N− depleted conditions, called low expression regions (LERs). Out of ten such regions, we have deleted two largest LERs, LER1 and LER2. The LER1 deletion (~110 kb deletion) and the LER1-LER2 serial deletion (~214 kb in total) showed essentially normal growth, lipid contents, fatty acid saturation levels and photosynthesis. These findings raise an exciting and new possibility to build a minimal genome in *Nannochloropsis*, which can serve as the chassis strain for customized production of biomolecules via further metabolic engineering.

## Results

### Selection of genomic regions for targeted deletion

To determine the “non-essential” regions under a particular condition, we analyzed transcriptomic datasets of *Nannochloropsis oceanica* IMET1 that we previously published under nitrogen repletion (N+) and nitrogen depletion (N−) conditions ((66, 67); also available in the NanDeSyn database at http://nandesyn.single-cell.cn/; (Gong et al., 2020)). We identified ten such regions, named LER (for <u>l</u>ow or no <u>e</u>xpression <u>r</u>egions), based on the threshold of mapped mRNA-Seq reads < 10 under N+ and N− conditions (average genome-wide sequence coverage of 53.3), i.e., LER1 through LER10 (**Table S1**). Among these, LER1 is the largest, and located at the distal end of Chr 30 (Fig. 1A). LER1 harbors 22 annotated genes (NO30G00010 - NO30G00220; details in NanDeSyn (78)) spanning over 98 kb of Chr 30 (**Table S1**). Among these, 4 genes encoded unknown functions, six membrane proteins, two PAS proteins, and so on (Table 1). These genes showed no or very low expression level under N+ or N− (Fig. 1B). Similar expression patterns of homologous genes are also found in the related strain *N. oceanica* CCMP1779 (45). Such a conserved low-expression pattern suggests that they are not essential for normal growth, at least under the nitrogen-related conditions. Interestingly, homologs of these genes was absent in *N. gaditana* B-31 and *N. salina* CCMP1776 (Fig. 1B; (44)), corroborating their non-essentiality in *N. oceanica*. Therefore, we hypothesized that this Chr 30 region is non-essential to *N. oceanica* and could likely be removed without compromising growth or other key traits.

**Table 1.** The genes annotation in the deleted region of Chr 30 (0-98305 bp).

| Gene ID | Annotation |
| --- | --- |
| NO30G00010 | zinc metalloendopeptidase, partial |
| NO30G00020 | unknown |
| NO30G00030 | hypothetical protein Naga_100716g2 |
| NO30G00040 | hypothetical protein SELMODRAFT (integral component of membrane) |
| NO30G00050 | unknown |
| NO30G00060 | unknown |
| NO30G00070 | hypothetical protein Naga_100209g9 |
| NO30G00080 | unknown |
| NO30G00090 | putative transmembrane protein (integral component of membrane) |
| NO30G00100 | putative transmembrane protein (integral component of membrane) |
| NO30G00110 | U6 snRNA-associated Sm-like protein Lsm3 (IPR040002) |
| NO30G00120 | hypothetical protein FisN_34Hu042 |
| NO30G00130 | hypothetical protein Naga_100380g2 (integral component of membrane) |
| NO30G00140 | hypothetical protein DRE_02121 (integral component of membrane) |
| NO30G00150 | hypothetical protein SD81_20880 |
| NO30G00160 | hypothetical protein Naga_100561g2 |
| NO30G00170 | hypothetical protein Naga_100561g2 |
| NO30G00180 | hypothetical protein Naga_100561g2 |
| NO30G00190 | collagen triple helix repeat protein |
| NO30G00200 | hypothetical protein Naga_100380g2 (integral component of membrane) |
| NO30G00210 | conserved unknown protein (with PAS-like domain ) |
| NO30G00220 | conserved unknown protein (with PAS-like domain ) |

**Figure 1.**
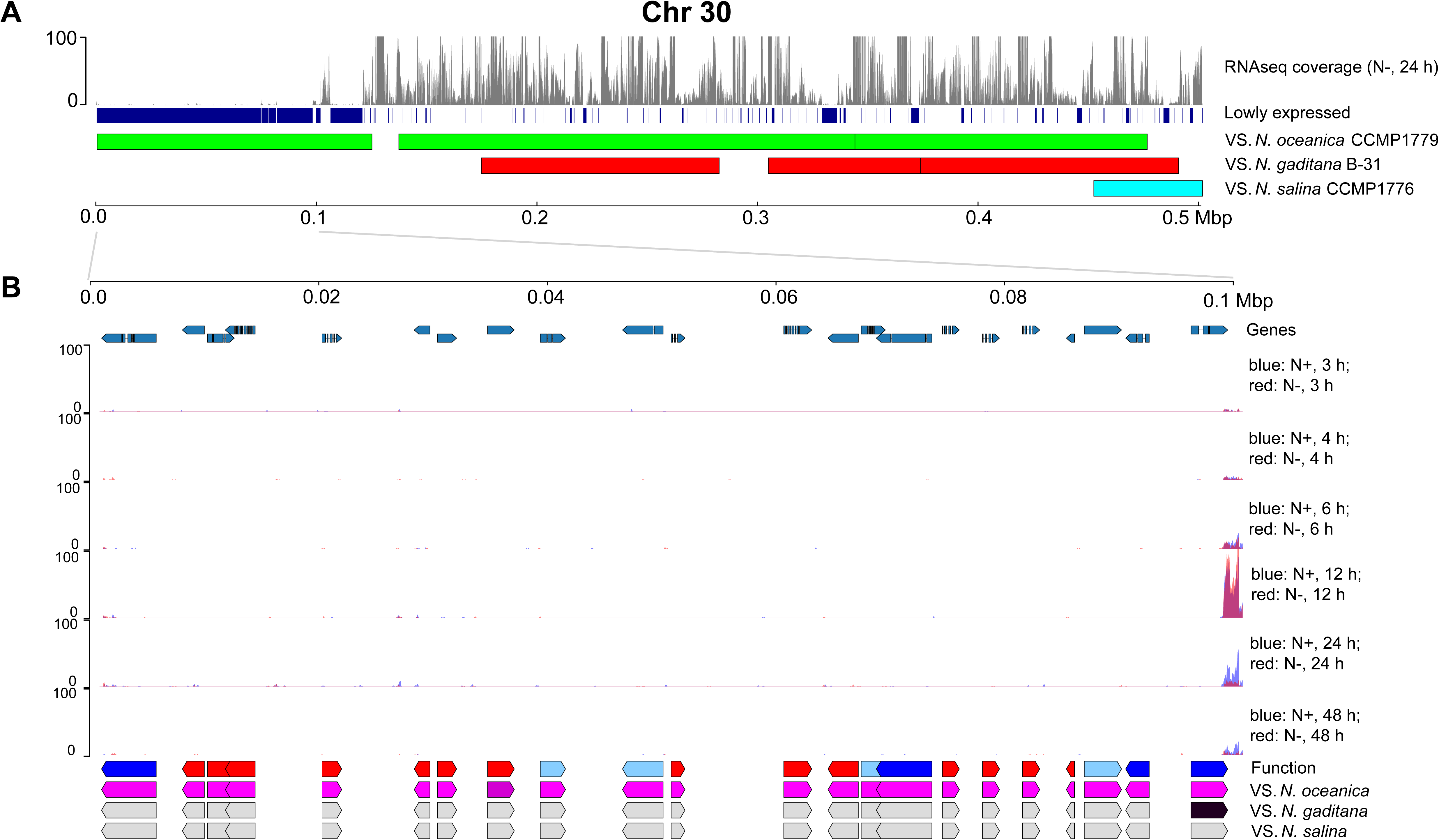
Rational selection of specific *N. oceanica* genomic regions targeted for deletion. (**A**) Transcriptome (N− 24 h) and genomic landscape of the Chr 30. Genomic fragments with low RNA-Seq expression (coverage < 10) were marked in blue. Synteny blocks between *N. oceanica* IMET1 and *N. oceanica* CCMP1779 (green), *N. gaditana* B-31 (red), *N. salina* CCMP1776 (light-blue) were also shown. (**B**) Transcriptome expression and potential function of 22 genes located in first 100 kb of the Chr 30 under N− and N+. For function row, genes with definite annotations were shown in blue; genes without any homologous genes were shown in red; genes with putative functions were shown in light-blue. The last three rows showed the existence of homologous genes (blastn, e-value < 1e-10) in *N. oceanica* CCMP179, *N. gaditana* B-31 and *N. salina* CCMP1776, with pink for existence, gray for non-existence.

### Deletion of LER1 and molecular validation of mutants

Episome-based CRISPR/Cas9 allowed removal of the circular extrachromosomal vector in the absence of selection pressure after stable mutagenesis was completed. Specifically, Cas9 and gRNAs were expressed under the endogenous ribosomal subunit bidirectional promoter (Pribi; Fig. 2A), which can drive dual expression of transgenes in *N. oceanica*. Two gRNAs were cloned for the deletion of LER1 (Fig. 2B,**Table S2**), and were separated by the hammerhead (HH) and hepatitis delta virus (HDV) self-cleaving ribozymes for their individual production (33). They were devised using chopchop (http://chopchop.cbu.uib.no/) (79), with gRNA1 located at ~20.5 kb distal to the telomere of Chr 30 (to avoid losing the telomere) and gRNA2 at ~81 kb proximal from gRNA1. Therefore, the two gRNAs were designed to cleave and delete ~81 kb inside from the Chr 30 telomere (Fig. 2B).

**Figure 2.**
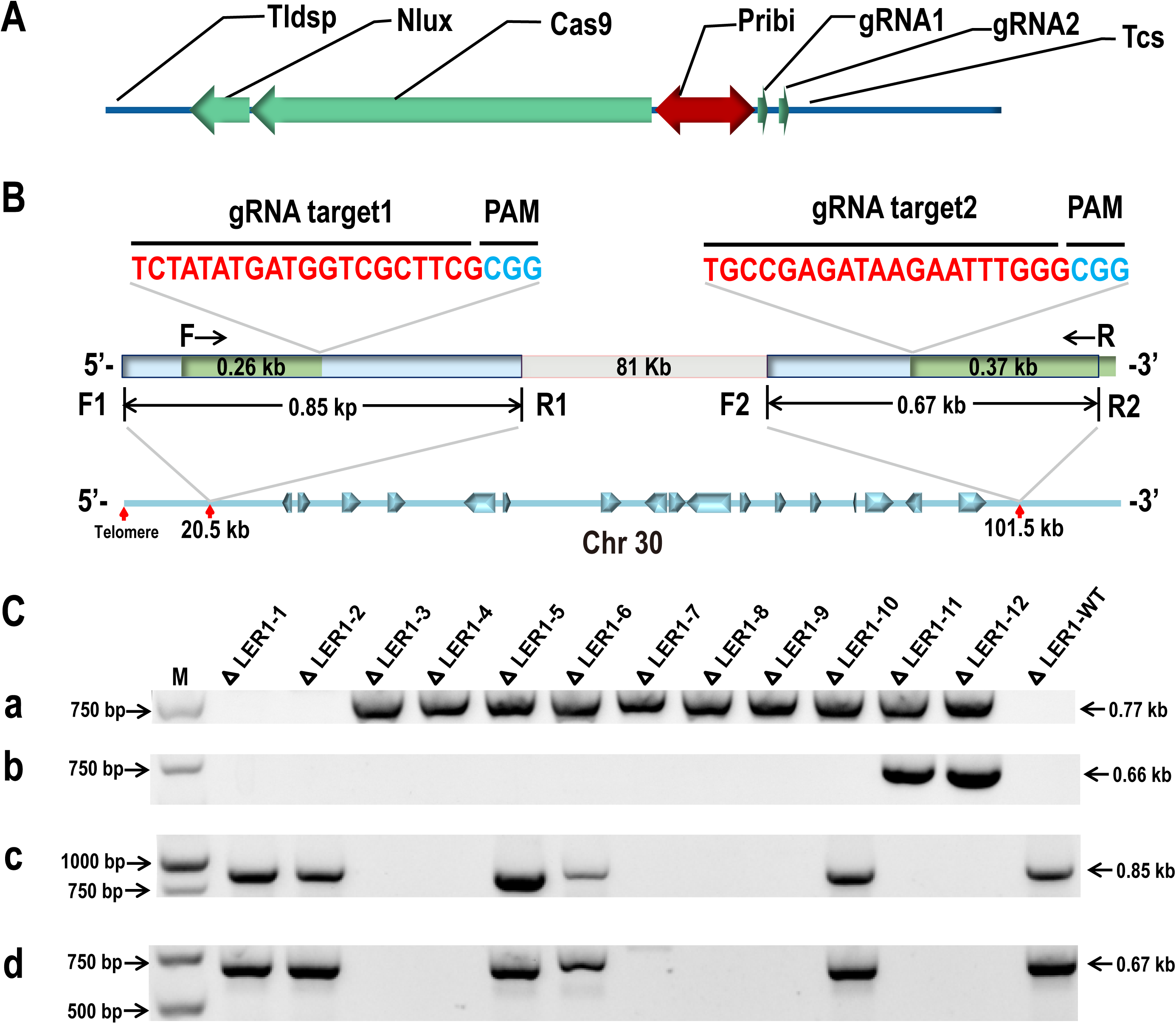
Vector design and PCR identification for Cas9/gRNA-mediated large fragment genome editing in IMET1. (**A**) Vector design for Cas9/gRNA medicated large fragment deletion in *Nannochloropsis oceanica* IMET1. The Cas9/gRNA constructs expressed two gRNAs and Cas9 from the Ribi promoter (Pribi). The gRNAs were cleaved by the HH and HDV ribozymes, once they were transcribed. (**B**) Design of gRNA target sites and target regions detection. The sites located at 20548 to 20567 and 101535 to 101554 of Chr 30, respectively, with 17 genes between them. PCR primers for the amplification of flanking region of target site 1 and target site 2 chromosomal deletions with F1 and R1, and F2 and R2, respectively. Deletion of the 81 kb internal fragment was confirmed by F and R primers. (C) Genomic DNA PCR results. (**a**) Gel image of the PCR products for detection of the plasmid-∆LER1. (**b**) Genomic DNA PCR for correct deletion between the cleavage sites of gRNA 1 and gRNA2. (**c, d**) Genomic DNA PCR for intact flanking regions around cleavage sites of gRNA1 and gRNA2, respectively. M, DNA marker.

We employed episome-based delivery of Cas9 and gRNAs, and the circular vector was initially maintained under selection pressure, which can be removed by non-selective media. The transformed plasmid-∆LER1 was selected on solid plates containing 300 μg/ml hygromycin and 1.6 g/L NaHCO3 for 25 days. Twelve colonies were cultured in selective liquid media, and their genomic DNAs were isolated and subjected to PCR for the presence of the plasmid-∆LER1 and chromosomal deletions (Fig. 2B, **Table S3**). Transformants 3-12 (∆LER1_3 to ∆LER1_12) were positive for the plasmid-∆LER1, while ∆LER1_1 and ∆LER1_2 were negative (Fig. 2C-a), suggesting that ∆LER1_3 to ∆LER1_12 were true transformants. For the genomic status of chromosomal deletions, only ∆LER1_11 and ∆LER1_12 showed amplification of 0.66 kb using primers F and R (Fig. 2C-b), suggesting correct deletion of the 81 kb between target sites cleaved by gRNA1 and gRNA2. We also checked for the status of flanking sequences around the cleavage site of gRNA1 (Fig. 2C-c) and gRNA2 (Fig. 2C-d) via primer pairs F1/R1 and F2/R2, respectively. ∆LER1_5, ∆LER1_6 and ∆LER1_10 were positive for these PCR reactions (similar to WT), suggesting that they contained cleavage sites of gRNA1 and gRNA2. Sanger sequencing of PCR products from T11 and T12 revealed correct deletion junction between the cleavage sites of gRNA1 and gRNA2, despite the presence of small indel mutations at the cleavage sites (Fig. 3A).

**Figure 3.**
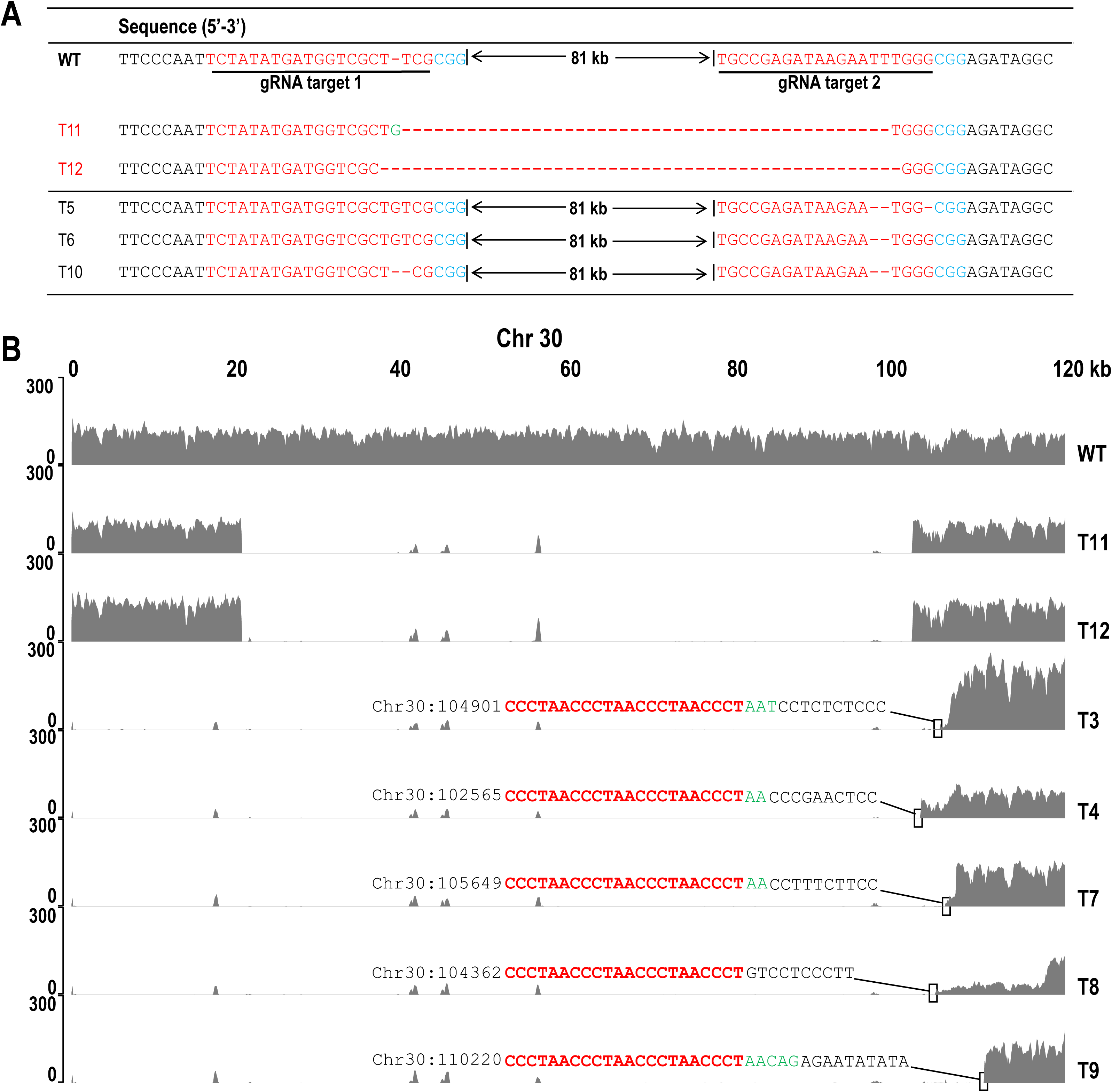
Genotypic validation of the mutants via both targeted and whole-genome shotgun sequencing. (**A**) Sanger sequence of the PCR products amplified from cleavage site 1, cleavage site 2 and the deletion in-between. (**B**) Summary of the whole genome sequencing of ∆LER1_3, ∆LER1_4, ∆LER1_7, ∆LER1_8, ∆LER1_9, ∆LER1_11 and ∆LER1_12 for their 5’ end of Chr 30. The new telomere was shown in red; genome sequence was shown in black; the indel was shown in green; the number indicates the original coordinate (in the WT chromosome) that corresponds to the first base of the newly formed terminal.

Interestingly, ∆LER1_3, ∆LER1_4, ∆LER1_7-∆LER1_9 were negative for all genomic PCR (Fig. 2C-b, c, d), suggesting that they lack all of the primer sites possibly by farther deletions on Chr 30. To confirm the exact nature of their chromosomal status, we sequenced their whole genome via NGS (Fig. 3B; **Methods**). We also sequenced the ∆LER1_11 and ∆LER1_12 genomes to probe whether they contained the distal part from the cleavage sites of Chr 30. The NGS data revealed that ∆LER1_11 and ∆LER1_12 contained correct distal sequences from the cleavage sites (Fig. 3B). However, ∆LER1_3, ∆LER1_4, ∆LER1_7-∆LER1_9 lacked not only the distal sequence of Chr 30 but also farther deletions (beyond the gRNA2 cleavage site) towards the 3’ ends of Chr 30 (Fig. 3B). Their endpoints towards the 3’ side varied in mutants, where up to 104900 bp were deleted for ∆LER1_3, 102564 bp for ∆LER1_4, 105648 bp for ∆LER1_7, 104361 bp for ∆LER1_8 and 110219 bp for ∆LER1_9. Therefore, ∆LER1_9 contained the largest deletion of Chr 30, where 110 kb was deleted (leaving only 392 kb as Chr 30).

For the extended deletion mutants (∆LER1_3, ∆LER1_4 and ∆LER1_7-9), we examined their 5’ termini, since telomeres are important for chromosome stability. To determine whether the ends maintained their own telomere or were replaced with new telomeres, we amplified the termini of these deletion mutants (**Table S4**) and cloned them into pXJ70gb (GenBank MT134322). Sequencing revealed the variable length of new ends to gRNA2 sequence in the range of 1.0 - 8.6 kb, which appeared as short CCCTAA repeats at the end of mutated Chr 30 (Fig. 3B), reminiscent of telomeric repeat structures in other organisms (80). Therefore, the telomere can regenerate randomly at the ends of chromosome in *N. oceanica*.

Finally, we probed the off-target effects in ∆LER1_3, ∆LER1_4, ∆LER1_7, ∆LER1_8, ∆LER1_9, ∆LER1_11 and ∆LER1_12. The potential off-target sites were predicted genome-wide for gRNA1 and gRNA2, by analyzing the assembled genome sequence of each of the mutants using Cas-OFFinder (81). Fourteen likely off-target sites within five nucleotide mismatches to the recognition site of gRNA1 (**Table S5**) were identified and 24 off-target sites were screened for gRNA2 (**Table S6**). Importantly, none of these sites were mutated in the transformants, as confirmed by their whole-genome sequences. The zero or very low off-target effects of CRISPR/Cas9 deletion mutants here encourage further rational genome-wide deletions with carefully selected gRNAs.

### Phenotypes of the LER1 deletion mutants

#### Growth, biomass, and photosynthesis

To probe the phenotype of these deletion mutations, we grouped mutants into (*i*) ∆LER1_11 and ∆LER1_12 with precise 81 kb deletions, and (*ii*) ∆LER1_3 (~104.9 kb removed), ∆LER1_4 (~102.6 kb removed) and ∆LER1_9 (the largest deletion of ~110.2 kb) with larger deletions. Under f/2 medium in flasks, shaking with 120 rpm under 40 μmol photos m-2s-1 at 23 °C, we measured their basic phenotypes including growth with OD750 (Fig. 4A and 4G), and biomass in μg/10ml (Fig. 4B and 4H). We also estimated photosynthetic activity by measuring Fv/Fm, as a ratio of variable to maximal fluorescence reflecting the optimal/maximal photochemical efficiency of PS II in the dark (82) and non-photochemical quenching (NPQ) which plays a major role in response to changes in light intensity in plants (83) (Fig. 4C and 4I). Overall, it was a surprise to find no or just subtle differences (e.g., Fv/Fm in T9 is 3% higher than that in WT; see below) in these phenotypes with such large chromosomal deletions.

**Figure 4.**
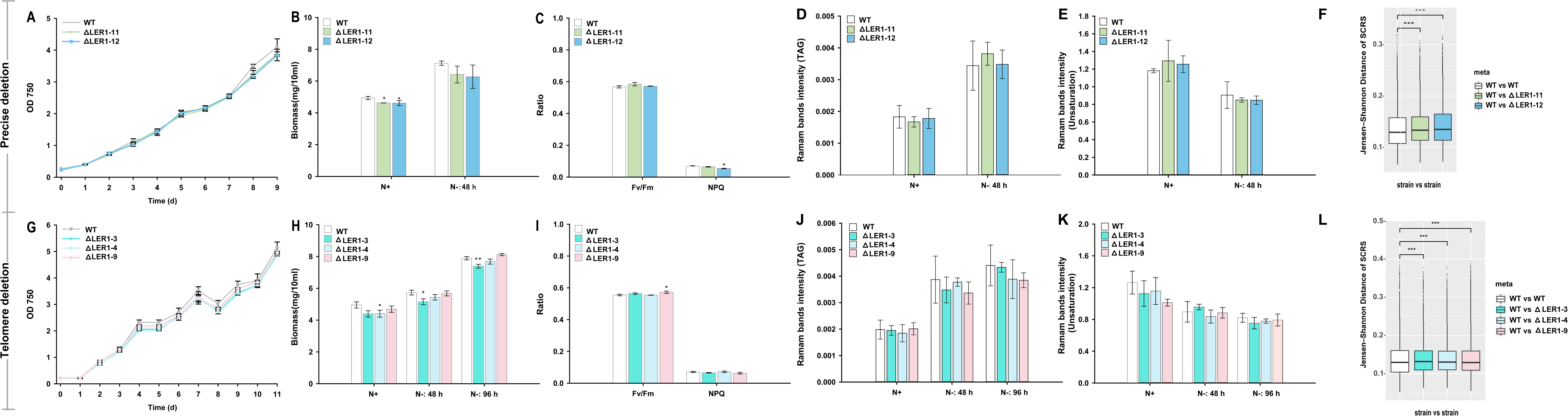
Phenotypic characterization of ∆LER1_3, ∆LER1_4, ∆LER1_9, e11 and ∆LER1_12. (**A**) The growth curve for ∆LER1_11, ∆LER1_12 and WT under N+ and N− 48 h. (**B**) The biomass for ∆LER1_11, ∆LER1_12 and WT under N+ and N− 48 h. (**C**) Optometric measurement of photosynthetic efficiency (Fv/Fm) and photoprotection in terms of NPQ for ∆LER1_11 and ∆LER1_12. (**D**) TAG content predicted by Raman band of 2851 cm^−1^ for ∆LER1_11, ∆LER1_12 and WT. (**E**) Degree of lipids unsaturation predicted by the ratio of 1656 cm^−1^ and 1640 cm^−1^ for ∆LER1_11, ∆LER1_12 and WT. (**F**) Comparison of inter-strain-Ramanome (WT-∆LER1_11 and WT-∆LER1_12) and intra-WT-Ramanome via Jensen-Shannon distance. (**G**) The growth curve for ∆LER1_3, ∆LER1_4, ∆LER1_9 and WT. (**H**) The biomass for ∆LER1_3, ∆LER1_4, ∆LER1_9 and WT under N+, N− 48h and N− 96h. (**I**) Photosynthetic efficiency (Fv/Fm) and NPQ for ∆LER1_3, ∆LER1_4 and ∆LER1_9. (**J**) TAG content predicted by Raman band of 2851 cm^−1^ for ∆LER1_3, ∆LER1_4, ∆LER1_9 and WT. (**K**) Lipids unsaturation degree predicted by the ratio of 1656 cm^−1^ and 1640 cm^−1^ for ∆LER1_3, ∆LER1_4, ∆LER1_9 and WT. (**L**) Comparison of inter-strain-Ramanome (WT-∆LER1_3, WT-∆LER1_4 and WT-∆LER1_9) and intra-WT-Ramanome via Jensen-Shannon distance. Pairwise Jensen-Shannon distances (JSD) of SCRS were calculated, and then JSD of inter-strain-Ramanome and intra-WT-Ramanome was stated. *: *p* value (t.test) <0.05; **: *p* value (t.test) <0.01;***: *p* value (wilcox.test) <0.001.

Growth measured via OD750 in the 81 kb-deletion mutants ∆LER1_11 and ∆LER1_12 was basically identical to WT (Fig. 4A), even though their biomass yield slightly decreased (Fig. 4B). Their photosynthetic parameters, Fv/Fm and NPQ, were identical, even though T12 showed moderate but significant reduction in NPQ (Fig. 4C). Larger-deletion mutants T3, T4 and T9 showed similar phenotypes compared to the 81 kb deletion mutants, in terms of growth (Fig. 4G) and biomass production (Fig. 4H). Photosynthetic parameters were also mostly equivalent, even though T9 showed slight but significant increase in the photosynthetic efficiency (Fig. 4I). These results suggest that genes included in the deleted areas of Chr 30 are not essential or critical for growth and photosynthesis under condition tested.

#### Lipid and degree of unsaturation in fatty acids via Ramanome

Via Single-cell Raman Spectra (SCRS), a ramanome can unveil single-cell-resolution phenomes in a label-free and non-invasive manner, e.g., characterize energy-storage molecules such as TAGs, starch and protein in *N. oceanica* (57, 67, 74, 84, 85). Therefore, to test whether the deletion mutants are phenotypically distinct from WT, we collected ramanome data of cells under N− condition (Batch1: WT/∆LER1_11/∆LER1_12 at 0 h, 48 h; Batch2: WT/∆LER1_3/∆LER1_4/∆LER1_9 at 0 h, 48 h, 72 h; **Methods**). TAG content as predicted by the intensity of Raman band of 2881 cm^−1^ showed no obvious difference between WT and mutants (Fig. 4D, J), and so was the degree of unsaturation (DU) for lipids, predicted by the ratio of 1656 cm^−1^ and 1640 cm^−1^ (Fig. 4E, K; (76)).

In addition to the TAG content and DU prediction, SCRS can also estimate the ‘fingerprint’ of a cell (86–88). Jensen-Shannon distances (JSD), which usually adapted for measuring the difference of frequency spectra (89, 90), could be used to measure the phenotypic difference between pairwise cells based on its SCRS. Moreover, to compare the phenotype difference among strains, here, we proposed ‘strain-Ramanome’ to define a certain strain, which consists of ramanomes of one certain strain at multiple inducing conditions and timepoints. For example, the ∆LER1_3-ramanome includes ramanomes of 0 h/ 48 h/ 72 h at N− of ∆LER1_3 transformants.

We calculated pairwise Jensen-Shannon distances (JSD) of intra-strain and inter-strain ramanomes based on JSD of the underlying SCRS (**Methods**). The results showed that JSDs of inter-WT-∆LER1_11/∆LER1_12/∆LER1_3/∆LER1_4/∆LER1_9 are significantly larger than intra-WT distance, which meant that ∆LER1_11/∆LER1_12/∆LER1_3/∆LER1_4/∆LER1_9 were all phenotypically different from WT strain (Wilcox test, *p* < 0.001, Fig. 6F, L). In conclusion, there is no apparent difference in TAG content or lipids unsaturation degree, while deletion mutants are phenotypically distinct from WT via strain-Ramanome analysis.

#### Temporal dynamics of transcriptome between WT and ∆LER1_9

To detect the gene-expression response of large fragment deletion, the transcriptomic profile of ∆LER1_9, the strain with the largest fragment (at 110 kb and harboring 24 genes) deleted among all mutants, were compared to that of WT by mRNA-Seq, over the three time points of 0 h, 48 h, 96 h (**Fig. S1**; **Methods**). In ∆LER1_9, the first 24 genes were deleted in Chr 30 and none of these transcripts was detected, although NO30G00230 was highly transcribed at each of the three timepoints in WT (which is consistent with the transcriptome data (67)). Thus, transcriptome results validated the large fragment deletion (Fig. 5A, **Fig. S2**).

**Figure 5.**
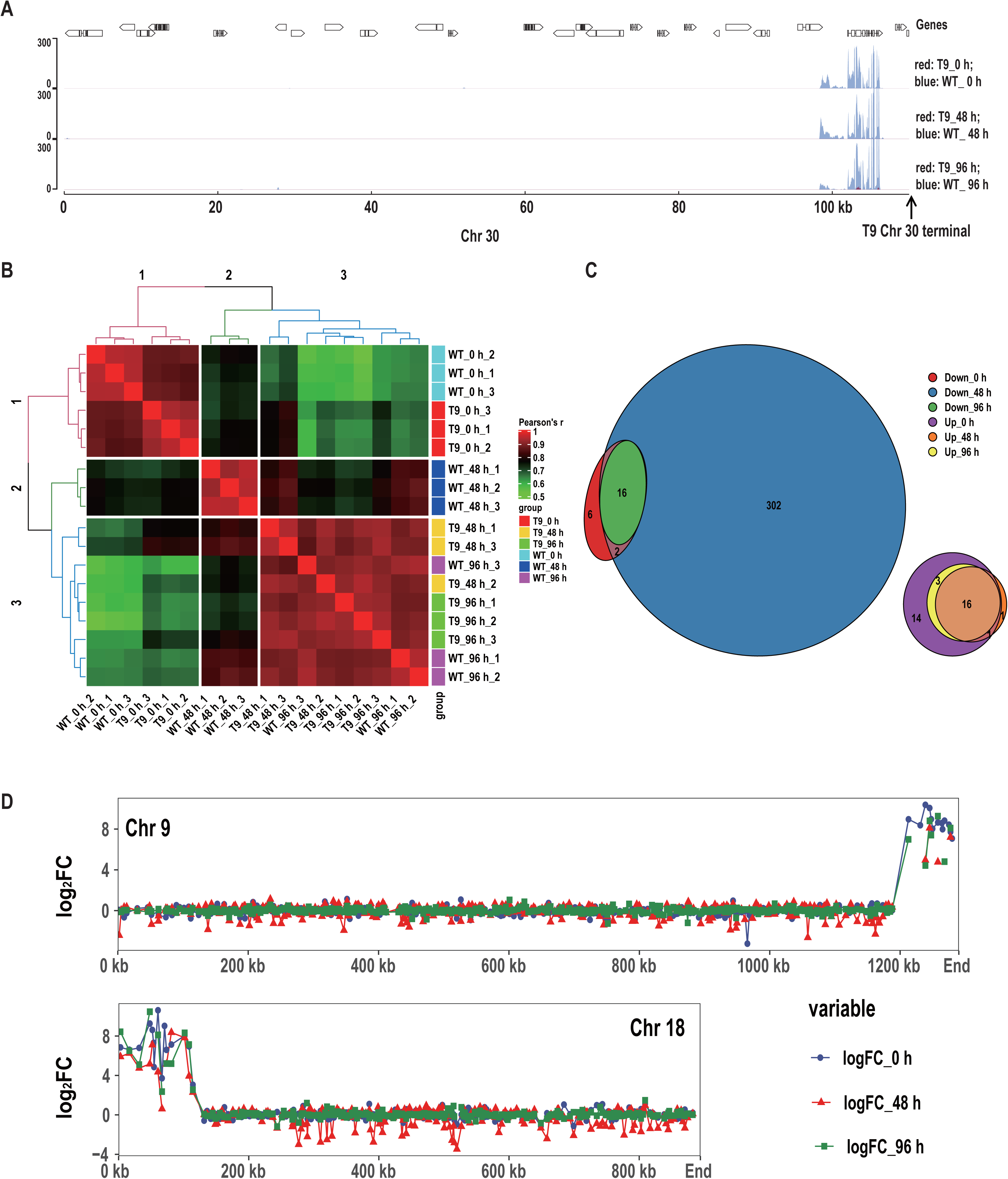
RNA-Seq analysis of ∆LER1_9 and WT. (**A**) RNA-Seq read mapping for ∆LER1_9 and WT under N+ 0 h, N− 48 h and N− 96 h. For ∆LER1_9, almost no reads were mapped to the 0-110000 region of Chr 30, suggesting the successful deletion of this large genomic region. (**B**) Clustered heatmap illustrating similarities of gene expression between different samples. Samples of ∆LER1_9 and WT were similar at N+ 0 h (Cluster 1, read branches) and N− 96 h (Cluster 3, blue branches). For N− 48 h, samples of ∆LER1_9 were similar to N− 96 h (all in Cluster 3), but samples of WT (N− 48 h) were still in an intermediate state (Cluster 2, green branches). (**C**) Venn diagram showing the numbers and overlap of differential expressed genes at 0 h, 48 h, and 96 h under N−. It’s remarkable that, under N− 48 h, 320 genes were down-regulated for ∆LER1_9 (compared to 24 genes under N+ and 16 genes under N− 96 h). (**D**) The differential expressed genes (at 0 h, 48 h, and 96 h under N−) are concentrated on the ends of Chr 9 and Chr 18. Besides, for N− 48 h, the remaining differential expressed genes spread over whole genome.

Correlation analysis of ∆LER1_9 and WT transcriptomes revealed two clusters: (*i*) ∆LER1_9 N-48 h, ∆LER1_9 N-96 h and WT N-96 h, and (*ii*) WT N+, ∆LER1_9 N+ and WT N-48 h. Compared to WT N-48 h, ∆LER1_9 N-48 h is more similar to ∆LER1_9 N-96 h and WT N-96 h. Thus ∆LER1_9 responds more quickly to N− than WT (Fig. 5B). Specifically, ~300 genes are down-regulated under N-48 h in T9 (vs. WT; Fig. 5C, **S3**), with no significantly-changed GO terms identified (**Fig. S4**).

Notably, in ∆LER1_9, 16 genes are upregulated under each of N+, N-48 h and N-96 h, as compared to WT (Fig. 5C), with most of those located near the ends of Chr 9 and Chr18 (Fig. 5D). Therefore, the large fragment deletion has likely changed chromosome conformation, resulting in the change of gene expression and leading to the rapid response of ∆LER1_9 under N−.

### Dual deletion of LER1 and LER2 in one round of transformation

To accelerate fragment deletion, we next tested the possibility to serially delete multiple fragments via one transformation with one vector. Four gRNAs were designed for the deletion of LER1 and LER2, which were separately cleaved by HH and HDV self-cleaving ribozymes for their individual production, respectively. Specifically, gRNA1 and gRNA2 were designed for LER1 deletion as described earlier (Fig. 6A-a; **Table S7**). LER2 was located at 1189260-1290044 in Chr 9, with very low gene expression (read depth<10). Therefore, gRNA3 were designed at ~100.7 kb (1189295 to 1189314) inside the 3’ telomere of Chr 9. To avoid losing the telomere, gRNA4 was designed to be located at ~19.5 kb (1270540 to 1270559) inside the 3’ telomere of Chr 9. Therefore, gRNA3 and gRNA4 were designed to cleave and delete ~81 kb inside from the telomere of Chr 9 (Fig. 6B-a; **Table S7**).

**Figure 6.**
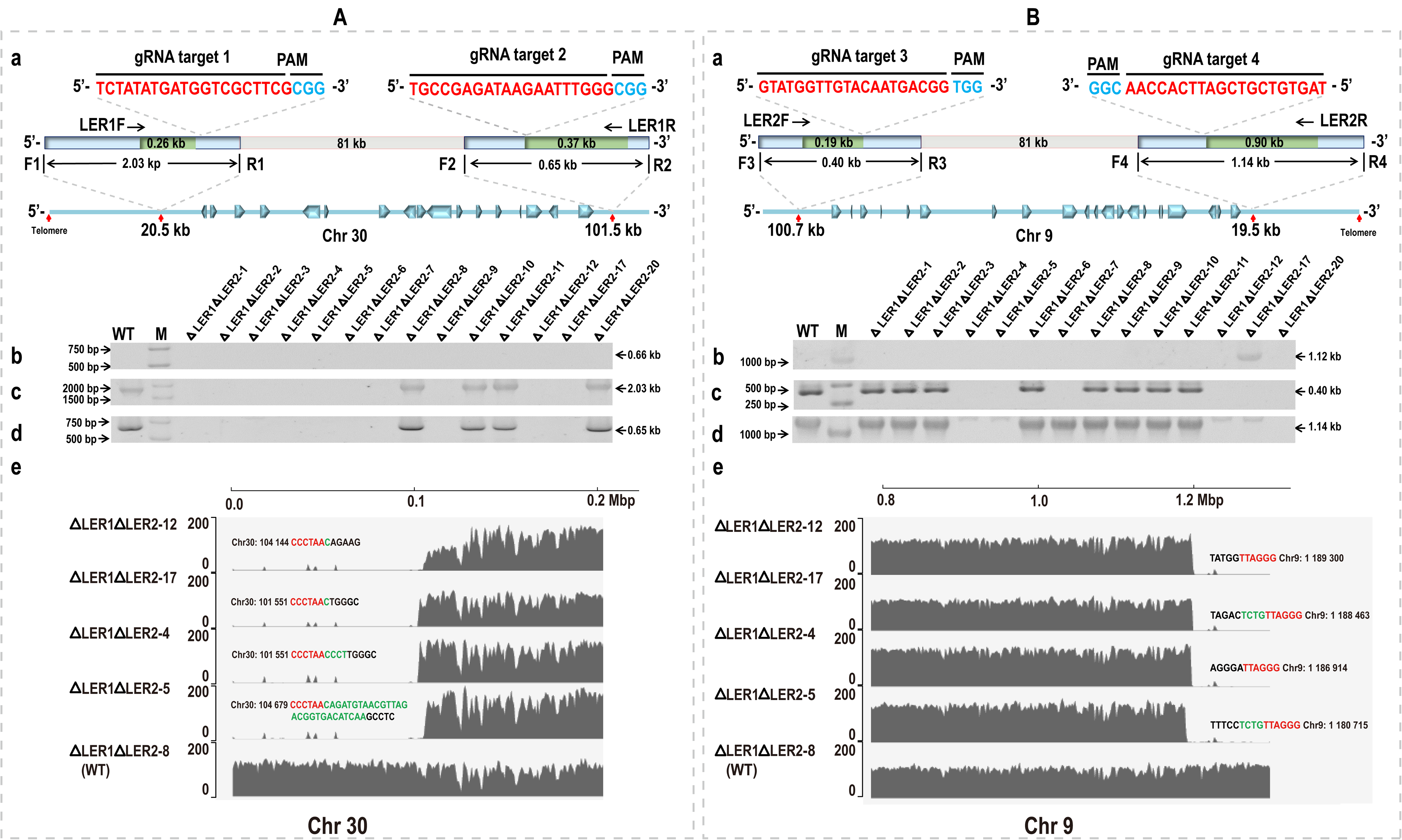
gRNAs design and transformant identification with genomic PCR and NGS for double large fragments deletion. (**A**) and (**B**) is the gRNAs design and transforamts identification with genomic PCR and NGS for LER1 deletion and LER2 deletion, respectively. (**a**) Design of gRNA target sites and target regions detection. The sites located at 20548 to 20567 (gRNA1), 101535 to 101554 (gRNA2) of Chr 30 and 1189295 to 1189314 (gRNA3), 1270540 to 1270559 (gRNA4) of Chr 9, respectively. PCR primers for the amplification of flanking region of target sites 1, target site 2, target site 3 and target site 4 chromosomal deletions with F1/ R1, F2/ R2, F3/R3 and F4/R4, respectively. Deletion of the 81kb internal fragments in Chr 30 and Chr 9 were confirmed by LER1F/ LER1R and LER2F/ LER2R, respectively. Target sites by gRNAs were marked on the chromosomes and the distances of gRNA1, gRNA2, gRNA3 and gRNA4 to the nearest telomeres were 20.5 kb, 101.5 kb, 100.7 kb and 19.5 kb, respectively. (**b)** Genomic DNA PCR for correct deletion between the cleavage sites. (**c, d**) Genomic DNA PCR for intact flanking regions around cleavage sites of gRNA1 and gRNA2 or gRNA3 and gRNA4, respectively. (**e**) Summary of the whole genome sequencing of ∆LER1∆LER2_12, ∆LER1∆LER2_17, ∆LER1∆LER2_4, ∆LER1∆LER2_5 and ∆LER1∆LER2_8 for their 5’ end of Chr 30 and 3’ end of Chr 9. The new telomere was shown in red; genome sequence was shown in black; the indel was shown in green; the number indicates the original coordinate (in the WT chromosome) that corresponds to the first base of the newly formed terminal.

Twenty colonies were cultured in selective liquid media, and their genomic DNAs were isolated and subjected to PCR for the presence of plasmid-∆LER1∆LER2 and chromosomal deletions (**Table S8**). The PCR identification was positive for ∆LER1∆LER2_1-∆LER1∆LER2_12, ∆LER1∆LER2_17 and ∆LER1∆LER2_20 (**data not shown**), suggesting that plasmid-∆LER1∆LER2 was successfully transformed. No transformants showed amplification of 0.66 kb at LER1 (Fig. 6A-b), suggesting either lack of target deletion or the complete loss of the LER1 region. As for LER2, ∆LER1∆LER2_17 showed 1.12 kb amplification (Fig. 6B-b), consistent with correct deletion of the 81 kb between target sites of gRNA3 and gRNA4.

To discriminate between precise targeted-region deletion and loss of chromosomal regions that extend beyond the targeted region, the flanking sequences around the target sites of gRNA1 (Fig. 6A-c), gRNA2 (Fig. 6A-d), gRNA3 (Fig. 6B-c) and gRNA4 (Fig. 6B-d) were assessed via primer pairs F1/R1, F2/R2, F3/R3 and F4/R4, respectively (**Table S8**). ∆LER1∆LER2_8, ∆LER1∆LER2_10 and ∆LER1∆LER2_11 were positive for these PCR reactions (similar to WT), suggesting that they contain the flanking sequence of cleavage sites of gRNA1, gRNA2, gRNA3 and gRNA4. ∆LER1∆LER2_20 were positive only at gRNA1 and gRNA2 PCR reactions, while ∆LER1∆LER2_1, ∆LER1∆LER2_2, ∆LER1∆LER2_3 and ∆LER1∆LER2_6 were positive only at gRNA3 and gRNA4 PCR reactions. Thus just one of the two targeted regions was deleted in each of these transformants. ∆LER1∆LER2_7 is positive only at gRNA4, which proved DNA sequence around gRNA4 was in the genome. ∆LER1∆LER2_4, ∆LER1∆LER2_5 and ∆LER1∆LER2_12 are negative for the above primers, suggesting that all of the regions were deleted.

To confirm deletions of LER1 and LER2 in ∆LER1∆LER2_4, ∆LER1∆LER2_5, ∆LER1∆LER2_12 and ∆LER1∆LER2_17, their genomes were profiled by NGS (∆LER1∆LER2_8 also sequenced as control). In ∆LER1∆LER2_4, ∆LER1∆LER2_5, ∆LER1∆LER2_12 and ∆LER1∆LER2_17, LER1 and LER2 were both deleted and telomeres regenerated at the newly generated terminals of chromosomes (Fig. 6A-e and 6B-e). The new chromosomal terminals correspond to the coordinates of 101551-104679 in Chr 30 and those of 1180715-1189300 in Chr 9, suggesting regeneration of new chromosomal terminals near the cleavage sites.

For deletion related to LER2 in ∆LER1∆LER2_17, PCR results indicated accurate deletion of the ~81 kb target region in LER2 (1189312-1270546 at the original coordinate of WT). In addition, sequences of the PCR products are consistent with predicted sequence derived from accurate deletion of the target fragment in LER2 as designed (**6B-b**). However, the NGS results supported <u>complete loss of the whole LER2</u> (the ~101 kb region from 1189300 to 1290044 at the original coordinate of WT) that extends beyond the original target region, as few NGS reads were mapped to the terminal 101 kb (from 1189300 to 1290044) in the 3’ of Chr 9 (Fig. 6B-e). However, two of the NGS reads were found that support the presence of the junction that corresponds to precise deletion of the ~81 kb target portion of LER2 (i.e., deleting the 1189312-1270546 region at the original coordinate of WT). Thus ∆LER1∆LER2_17 is genetically heterogeneous, with majority of the cells being the complete deletion of the whole LER2 (~101 kb deleted; 1189300-1290044 at the original coordinate) and ~2% being the 81 kb precise deletion of the target region (1189312-1270546 region at the original coordinate of WT) in LER2. Therefore, altogether, there is evidence for successful deletion of both LER1 and LER2 in 4 of the 14 transformants, which validated the method for serial large-fragment deletion in *N. oceanica*.

### Phenotypes of the ∆LER1∆LER2 mutants with double deletion

To probe phenotypes of the double-deletion mutations, growth, biomass and photosynthesis were analyzed for in ∆LER1∆LER2_4, ∆LER1∆LER2_5 and ∆LER1∆LER2_17 under f/2 cultured for 7days (N+), N-48 h and N-96 h, as described earlier for LER1. Growth and biomass of these mutants were slightly elevated as compared to WT (Fig. 7A and B). For growth, under N+, ∆LER1∆LER2_17 increased by 10.7%, and under N-96 h, ∆LER1∆LER2_4 and ∆LER1∆LER2_17 increased by 10.1% and 4.8%, respectively. For biomass, ∆LER1∆LER2_4 increased by 10.4% under N+, and ∆LER1∆LER2_4 and ∆LER1∆LER2_17 increased by 12.0% and 5.1%, under N-96 h, respectively. Their photosynthetic parameters, Fv/Fm and NPQ, were mostly unaffected, even though T4 and T17 showed moderate but significant reduction in Fv/Fm (Fig. 7C). These results suggested fragment deletions is feasible to remold *N. oceanica* as chassis cell without affection of growth under specific environmental conditions.

**Figure 7.**
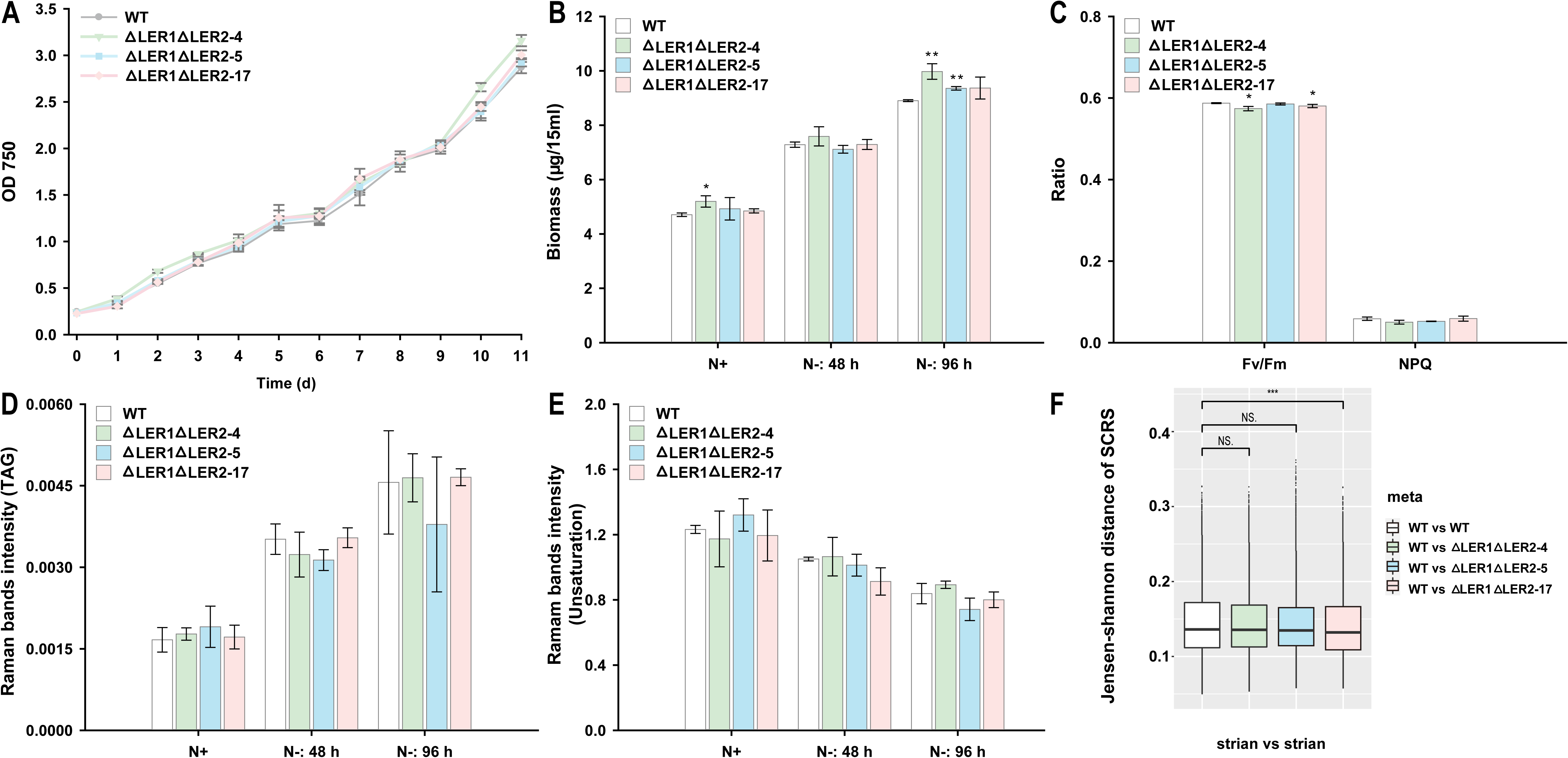
Phenotypic characterization of ∆LER1∆LER2_4, ∆LER1∆LER2_5, ∆LER1∆LER2_17 and WT. (**A**) The growth curve under f/2 medium cultured for 7 days and N− induced for 48 h and 96 h. (**B**) The biomass for ∆LER1∆LER2_4, ∆LER1∆LER2_5, ∆LER1∆LER2_17 and WT. (**C**) Optometric measurement of photosynthetic efficiency (Fv/Fm) and photoprotection in terms of NPQ for ∆LER1∆LER2_4, ∆LER1∆LER2_5, ∆LER1∆LER2_17 and WT. (**D**) TAG content predicted by Raman band of 2851 cm^−1^ for ∆LER1∆LER2_4, ∆LER1∆LER2_5, ∆LER1∆LER2_17 and WT (**E**) Degree of lipids unsaturation predicted by the ratio of 1656 cm^−1^ and 1640 cm^−1^ for ∆LER1∆LER2_4, ∆LER1∆LER2_5, ∆LER1∆LER2_17 and WT. *: *p* value (t.test) <0.05; **: *p* value (t.test) <0.01. (**F**) Comparison of inter-strain (WT vs. ∆LER1∆LER2_4; WT vs. ∆LER1∆LER2_5; WT vs. ∆LER1∆LER2_17) and intra-strain similarity of the ramanomes via the Jensen-Shannon distance. Pairwise JSDs of SCRS were calculated. ***: *p* value (wilcox.test) <0.001.

To test whether the TAG content and DU of deletion mutants are distinct from WT, ramanome data of ∆LER1∆LER2_4, ∆LER1∆LER2_5, ∆LER1∆LER2_17 and WT were collected under N− condition for 0 h, 48 h and 72 h. TAG content and DU was predicted as described earlier for LER1, and no apparent difference in TAG content or DU was detected between WT and mutants (Fig. 7D and E). JSD also was used to measure the phenotypic difference between mutants and WT based on its SCRS. The results showed that JSD of inter-WT/∆LER1∆LER2_17 were significantly larger than intra-WT distance, which meant that ∆LER1∆LER2_17 were phenotypically different from WT strain (Wilcox test, *p* <0.001, Fig. 7F). However, JSD of inter-WT/∆LER1∆LER2_4/∆LER1∆LER2_5 distance were similar with intra-WT distance, indicating no significant different between ∆LER1∆LER2_4/∆LER1∆LER2_5 and WT strain. In conclusion, there is no apparent difference in TAG content or lipids unsaturation degree, while dual-deletion mutant ∆LER1∆LER2_17 is phenotypically distinct from WT via strain-Ramanome analysis. These results demonstrated that large-fragment deletions, i.e., removal of both LER1 and LER2, exerts no effects on the *N. oceanica* phenotypes.

## Discussion

Microalgae have great potential as the next-generation feedstock for biofuels and chemicals in an eco-friendly manner; however, for most microalgae, the genetic toolboxes have been lagging behind crop plants and other microorganisms partly due to late development and inherent technical difficulties (91). This has hindered the exploitation of the extensive microalgal genomic resources for crafting a minimal microalgal genome with uncompromised functionality that can serve as a solar-energy driven, CO_2_-fixing chassis for green biomanufacturing. In animal and plants, the CRISPR system can produce deletions ranging between several hundred bp and a few hundred kb (92–95), however, target deletion of chromosomal regions has not been demonstrated in microalgae, one of the most diverse groups of organisms on Earth. In fact, whether and to what extent microalgal genomes can be molded is unknown.

To tackle this challenge, we exploited the rich functional genomic resources of *N. oceanica* to identify the “dispensable” chromosomal regions for targeted deletion, and also took advantage of its highly efficient DNA transformation system to generate deletion of designated chromosomal regions via CRISPR/Cas9. Specifically, bidirectional promoters were newly developed for both Cas9 and the dual gRNAs, and individual gRNAs were separately produced by ribozymes, all of which contained in a plasmid. We demonstrated one-time deletion of up to ~214 kb from *N. oceanica* Chr 30, which is 973 times longer than the genome fragments deleted in microalgae previously reported (the removal of 220 bp in *E. gracilis* genome using ribonucleoprotein or RNP (40)). Notably, our episome-based genome engineering did not leave any traces of foreign DNA, potentially avoiding the GMO conflict in the future (33, 96).

In the LER1 fragment deletion, among 10 positive transformants, 7 transformants deleted LER1 fragment. In LER1 and LER2 dual fragment deletions, among 14 positive transformants, 4 transformants deleted LER1 and LER2 by one transformation. These results indicated the deletion efficiency is enough to construct minimal genome. However, only 2/7 transforamnts with LER1 fragment deletion is accurate deletion and no accurate deletion transformants were found among LER1 and LER2 dual fragment deletions transformants. We speculated that the losing of untargeted sequence near the telomere is for its “unimportant”. If more accurate deletion transformants required, more transformants need to be screened.

Moreover, fidelity of our Cas9-mediated deletion of chromosomal segments is very high, since no off-targeting events were detected based on Cas-OFFinder (81) and whole genome sequencing for the deletion mutants (**Tables S5** and **S6**). However, even though we successfully obtained two mutants (∆LER1_11 and ∆LER1_12) with precise deletions at gRNA1 and gRNA2, a number of mutants contained deletions beyond the cleavage sites by gRNAs, e.g., in ∆LER1_3, ∆LER1_4, ∆LER1_7, ∆LER1_8 and ∆LER1_9, the whole distal segments were deleted (Fig. 3B), probably due to failure of correct ligation between the two cleavage sites by gRNA1 and gRNA2. Interestingly, sequencing of the deleted ends (Fig. 3B) revealed that the telomere appeared to be regenerated, containing repeats of CCCTAA which are typical telomeric repeats found in other organisms (80, 97, 98). The de *novo* addition of telomere to the end of DSBs protected the Chr 30 from shortening and maintained stability of the whole genome, as reported in yeasts (99). While mechanisms of *Nannochloropsis* telomere maintenance is unknown, our accidental discovery of autonomous telomere regeneration in *N. oceanica* is important, as this would guide artificial chromosomes construction (100) and telomere-mediated chromosomal truncation (101) in this and related organisms, and greatly expand the scope of genome engineering in microalgae. Notably, although their transcript level of the genes in LER1 at Chr 30 was low at f/2 medium under both N+ and N− conditions, a few were induced by the high CO_2_ level (50,000 ppm) (68). While no functional links with carbon metabolic pathways are apparent (Table 1), the LER1-harboring genes might be important to *N. oceanica* under other conditions. Nevertheless, they seemed to be unrelated to nitrogen-related metabolic pathways, consistent with our results showing only minimal or no phenotype change in growth, photosynthesis and lipids in the deletion mutants. Thus it would be encouraging to continue deleting other regions of the genome until we achieve the minimal yet functional *N. oceanica* genome, which can then be employed as a solar-energy driven, CO_2_-fixing chassis for green biomanufacturing.

## Materials and Methods

### Genome-wide screening/selection of candidate regions for genomic-region deletion

To selectthe LERs, we scanned the whole genome and N+/N− transcriptome of *N. oceanica* IMET1 from the NanDeSyn database (http://nandesyn.single-cell.cn). Low-expression genomic regions (local coverage <10) were detected using transcriptome dataset SRP017310 (67). Synteny blocks between different *Nannochloropsis* species were retrieved from NanDeSyn website (http://nandesyn.single-cell.cn/synview/search). Comparison of the gene expression of all chromosomes revealed a 98 kb genomic fragment with almost no genes expression in the beginning of Chr 30. Potential functions of genes within this fragment were manually checked according to gene feature pages on NanDeSyn website (e.g. http://nandesyn.single-cell.cn/feature/gene/NO30G00150). Multi-omics information was visualized using genome browser deposited in NanDeSyn website (http://nandesyn.single-cell.cn/browser) or plotted using pyGenomeTracks package (102).

### Construction of CRISPR/Cas expression vectors

An episome-based CRISPR/Cas system (pNOC-ARS-CRISPR-v2) was employed in this study (48). A pair of gRNAs were designed (**Table S2**) with the distance about 81 kb in Chr 30 at the positions of 20548 to 20567 and 101535 to 101554, respectively. The two gRNAs were expressed in one RNA molecule promoted by Pribi, and each of the gRNA was flanked by HH and HDV to allow precise cleavage. For the dual-fragment deletion, gRNA3 (1189295 to 1189314) and gRNA4 (1270540 to 1270559) that target Chr 30 were expressed together with gRNA1 and gRNA2 that targets Chr 9, via one single vector.

### Microalgal culture growth and transformation

*N. oceanica* strain IMET1 was maintained in the dark on solid f/2 medium (67), which contains 15 g/liter agar and 1.6 g/L NaHCO3 at 4 °C. For use in transformation experiments, cells were inoculated into liquid cultures of the f/2 medium and maintained under light at 50 μmol photos m-2 s-1 at 23 °C. The episome with CRISPR/Cas system was transformed into *N. oceanica* using the electrophoresis protocol we previously described (26). The growth curve was detected with f/2 medium in flasks, shaking with 120 rpm under 40 μmol photos m-2s-1 at 23 °C.

### Validation of the transformants with large fragment deletion

The genomic DNA of transgenic and wild-type *N. oceanica* cells was extracted. Episome PF and Episome PR were used to amplify the extracted DNA to detect the existence of episome. Primer F and Primer R were designed to detect the large fragment deletion in transformants. Primer F1, R1 and Primer F2, R2 were designed to amplify the gRNA target site 1 and gRNA target site 2, respectively (**Table S3**). PCR products were detected with Sanger sequencing to obtain the mutation sequence of the target sites. Moreover, to detect the terminal of ∆LER1_3, ∆LER1_4, ∆LER1_7, ∆LER1_8, ∆LER1_9, primers were designed according to the NGS results and telomere sequence. PCR products were ligated to the *Kpn* I digested pXJ70gb (GenBank MT134322), and the clones were sequenced with the Sanger method.

### Genome-wide mutation mapping of the transformants for detecting deletion events and potential off-target sequences

The genomic DNA was extracted with HP DNA Kit (Omega Bio-Tek, America). DNA was sheared to 300 bp and sequencing libraries were deep sequenced on Illumina Hiseq platform. Whole-genome sequencing libraries of eight samples were prepared using standard protocols for the Illumina HiSeq 4000 platform, generating about 3 gigabytes of raw data for each sample. The Illumina raw reads were trimmed using TrimGalore to remove adaptors and bases of low quality. Then, the cleaned reads were mapped to the reference genome from NanDeSyn database (http://nandesyn.single-cell.cn; IMET1v2) using the BWA mem program (103), resulting BAM files were visualized using Jbrowse genome browser (http://nandesyn.single-cell.cn/jbrowse). Clean reads were assembled using SPAdes (104) in multi-cell mode, with parameters to automatically compute coverage threshold (“--cov-cutoff auto”). Sequence variants were called for all samples using GATK. Variant calling and filtration using GATK software were performed with the HaplotypeCaller and VariantFiltration commands, respectively. The WGS data used in this study can be accessed at NanDeSyn database (ftp://nandesyn.single-cell.cn/pub/tracks/).

To identify any potential off-target sequences in the whole-genome sequence of Cas9/gRNA transformants, Cas-OFFinder (81) was used to find potential gRNA-DNA mismatch pairs in the whole genome where mismatched bases in each pair are less than or equal to 5. The 38 potential off-target sites were manually checked based on WGS data (variant calling results and reads alignment visualization).

To further characterize the deletion events, a pair of primers was designed to amplify the terminals (Primer F based on the sequence of telomere and primer R based on the NGS results; **Table S4**). The PCR products were cloned into pXJ70gb (GenBank MT134322), and subjected to Sanger sequencing.

### Photosynthesis parameter monitoring

Cells were grown in culture flasks for 4 days to the exponential phase on a shaker (125 rpm/min) at 23 °C under continuous light (40 μmol photons•m-2•s −1). Chlorophyll fluorescence of WT and mutants were measured using a pulse-amplitude modulated fluorometer (Image PAM, Walz, Effeltrich, Germany) after 20-min dark treatment of cells. PSII maximum quantum yield (Fv/Fm) and NPQ were measured according to a previous report (59, 105).

### Ramanome-based phenotyping of the transformants

Cell aliquots were collected right before re-inoculation (i.e. 0 h), and from each triplicate of Group N−: 48 h, and 96 h. Before measurement, each cell sample was washed three times and resuspended in ddH_2_O to remove the culture media, and then loaded in a capillary tube (50 mm length × 1 mm width × 0.1 mm height, Camlab, UK). Raman spectra of individual cells were acquired using a Raman Activated Cell Sorting system (RACS, Wellsens Inc, China), which was equipped with a confocal microscope with a 50 × PL magnifying dry objective (NA=0.55, BX41, Olympus, UK) and a 532 nm Nd:YAG laser (Ventus, Laser Quantum Ltd, UK). The power out of the objective was 200 mW and the acquisition time was 2 seconds per cell. Each Raman spectrum was acquired between the range 3340.9036 cm^−1^ and 394.11472 cm^−1^. About 20 cells were measured in each of the samples. For background spectrum, the average of three spectra acquired from the liquid around the cell was used.

Pre-processing of raw spectra was performed with LabSpec 6 (HORIBA Scientific), including background subtraction and the baseline correction by a polynomial algorithm with degree of seven. The whole spectra were normalized for further analyses. Moreover, pairwise Jensen-Shannon distances (JSD) of single-cell Raman spectrum (SCRS) were first calculated, and then JSD of inter-strain-Ramanome and intra-WT-Ramanome was derived.

### Transcriptome sampling, sequencing and analysis

To compare the temporal dynamics of transcriptome between the mutants and WT, ∆LER1_9 and WT were cultured in f/2 medium for 7 days and induced with N-deplete f/2 medium for 4 days. The samples were collected at 7 days (N-0 h), N-48 h, N-96 h. *N. oceanica* cells were harvested by centrifugation for 5 min at 2500 g and then were immediately quenched with liquid N2 and stored in − 80 °C freezer. Total algal RNA was extracted using Trizol reagents (Tiangen, Beijing, China). The concentration and purity of the RNA were determined spectrophotometrically (IMPLEN, CA, USA) and RNA integrity was assessed using the RNA Nano 6000 Assay Kit of the Agilent Bioanalyzer 2100 system (Agilent Technologies, CA, USA). A total amount of 2 μg RNA per sample was used as input material for the RNA sample preparations. Sequencing libraries were generated using NEBNext Ultra™ RNA Library Prep Kit for Illumina (NEB, USA) following manufacturer’s recommendations and index codes were added to attribute sequences to each sample.

The clustering of the index-coded samples was performed on a cBot Cluster Generation System using the HiSeq 3000/4000 PE Cluster Kit Box1 from Illumina. After cluster generation, the library preparations were sequenced on an Illumina HiSeq 4000 platform and 150bp paired-end reads were generated. Raw data (raw reads) of fastq format were quality controlled, aligned to the reference genome (IMET1v2) and generated gene abundances using nfcore/rnaseq pipeline (v1.2; https://doi.org/10.5281/zenodo.1400710).

Scripts bundled with Trinity software v2.4.0 (106) were mainly adopted to normalize gene abundances and find the differentially expressed subset. A table of TMM-normalized TPM expression matrix and a separate table of raw fragment counts were generated for further analysis and visualization. Differentially expressed (DE) genes were identified from raw counts with the Bioconductor package EdgeR v.3.16.5 (107). Three biological replicates for each condition were provided. The most significant differentially expressed genes (FDR < 0.001 and FC > 4) were extracted for further analysis. A hierarchically clustered heatmap was generated from the Pearson correlation matrix of pairwise sample comparisons based on the most significant DE subset.

## Author contribution

Q.W. and J.X. conceived the project. Y.G. conducted bioinformatics analysis. Y.H. performed Ramanome analysis. Q.W., Y.X., N.L., X.D., Y.L. generated mutants. Q.W. phenotypically characterized mutants. Q.W., J.X., BrJ interpreted phenotypic data. Q. W., Y.G. and J.X. analyzed transcriptomes of mutant and wild-type strains. J.X., Q.W., Y.G., BrJ and Y.H. wrote the paper.

## Acknowledgement

The work was supported by National Key Research and Development Program (2018YFA0902500 for J.X., Q.W. and Y.X.), Natural Science Foundation of China (31425002 for J.X.; 31800071 for Q.W.) and Chinese Academy of Sciences President’s International Fellowship Initiative (2020VBA0032 for BrJ).

