## Supplement Figure S1-S4 for "Genome engineering of *Nannochloropsis* with large deletions for constructing microalgal minigenomes"

**Figure S1. The scheme of N-depletion induction for ΔLER1_9 and WT.** ΔLER1_9 and WT cultured in f/2 medium for 7 days and induced with N- f/2 medium for 48 h and 96 h (3 replicates of each).

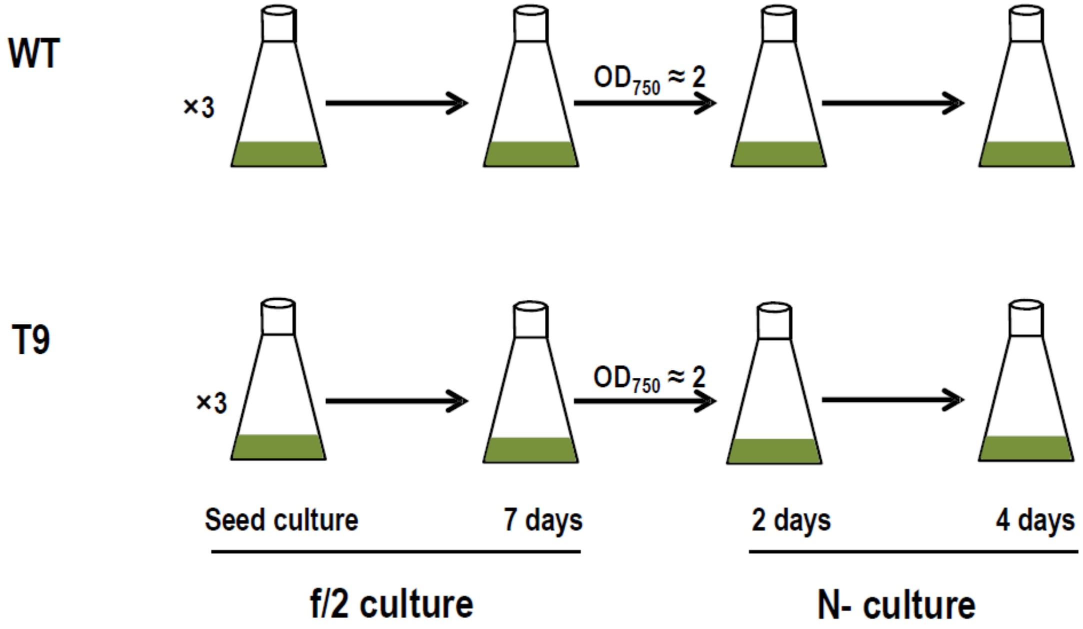

**Figure S2. Gene expression in the LER1 region in the ΔLER1_9 and WT lines.**

RNA-Seq reads mapped at the genomic region of chr30:97000-107000 for ΔLER1_9 and WT under N+ 0 h, N- 48 h and N- 96 h were shown. For ΔLER1_9, almost no reads are mapped to this genomic region, implying the successful deletion of the target region.

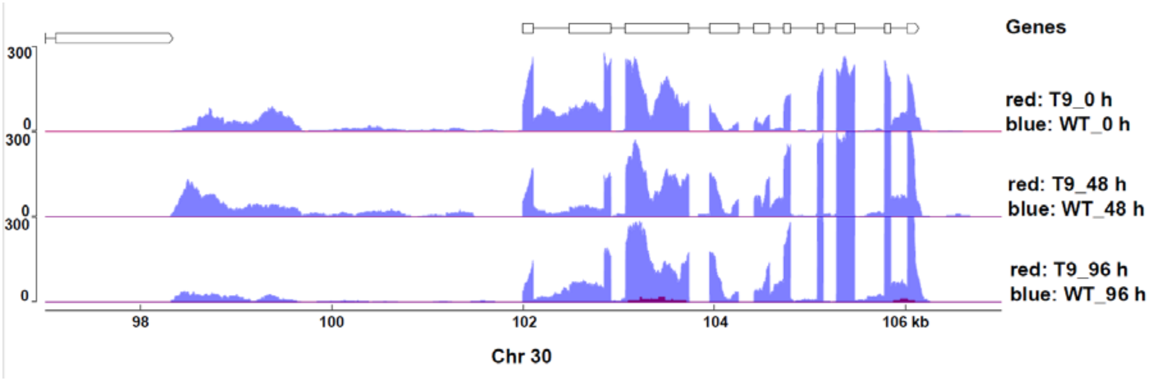

**Figure S3. Clustering analysis of the expression patterns of differentially expressed genes between the ΔLER1_9 and WT lines.** Heatmap of the clusters of differential expressed genes (or samples) was shown for ΔLER1_9 and WT (under N+ 0 h, N- 48 h and N- 96 h). Notably, N- 48 h and N- 96 h of ΔLER1_9 and N-96 h of WT were clustered together, suggesting a "quicker" response for ΔLER1_9 under N- stress.

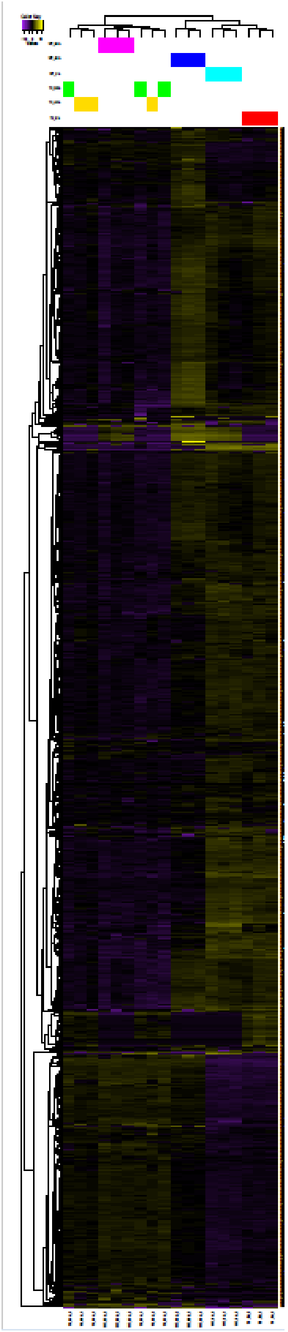

**Figure S4. GO-term analysis of the differentially expressed genes between the ΔLER1_9 and WT lines.** Distribution of GO terms is shown for differential expressed genes at N+ 0 h, N- 48 h and N- 96 h. Differential expressed genes were obtained by comparing between ΔLER1_9 and WT.

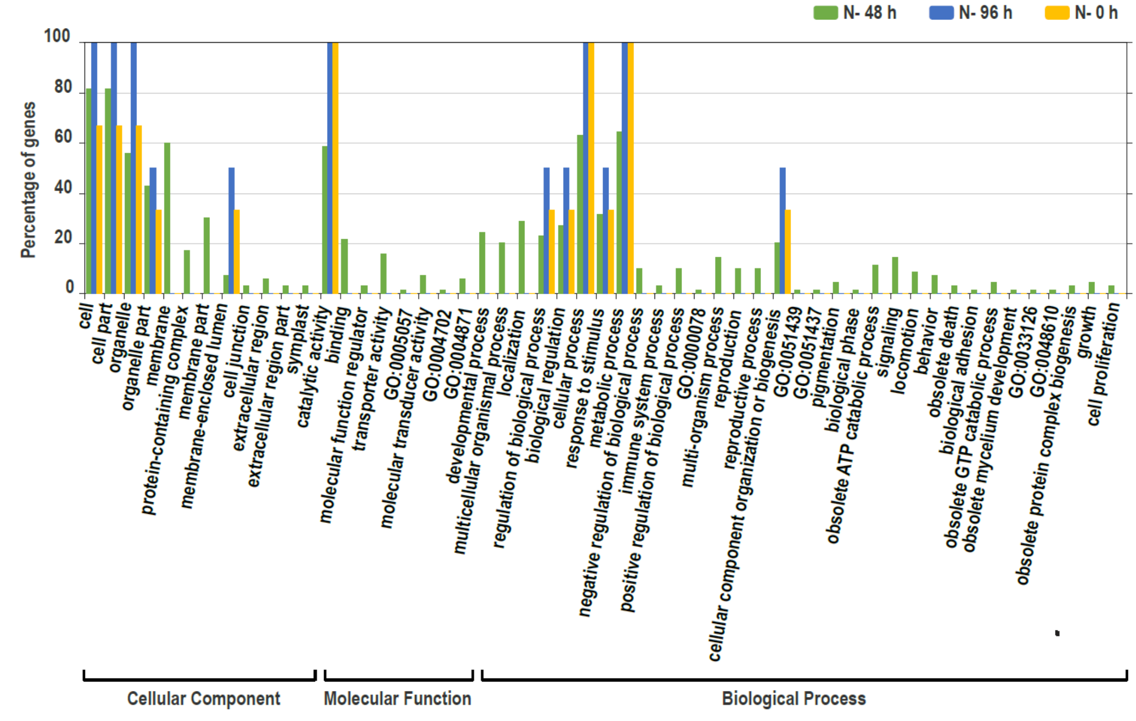

**Table S1. The list of *N. oceanica* genomic regions with minimal transcription of mRNA under N- and N+ conditions.** Genomic regions with fewer than ten mRNA-seq reads (except LER2 with read depth ≥10 around 1220423 under N+ 24 h) from dataset of SRP017310 were searched along chromosomes, and ordered by fragment lengths.

| Chromosome | Region | Fragment size (bp) | Genomic regions with minimal transcription |
|---|---|---|---|
| **Chr 30** | 0-98305 | 98305 | LER1 (deleted) |
| **Chr 9** | 1189260-1290044 | 100784 | LER2 (deleted) |
| **Chr 9** | 1220423-1290044 | 69621 | LER2’ (deleted) |
| **Chr 1** | 1445632-1491336 | 45704 | LER3 |
| **Chr 3** | 235377-278972 | 43595 | LER4 |
| **Chr 1** | 3661-40511 | 36850 | LER5 |
| **Chr 15** | 902831-933118 | 30287 | LER6 |
| **Chr 18** | 81714-111725 | 30011 | LER7 |
| **Chr 18** | 22858-47078 | 24220 | LER8 |
| **Chr 4** | 645188-668274 | 23086 | LER9 |
| **Chr 17** | 304475-326132 | 21657 | LER10 |

**Table S2. The gRNA target sequence for LER1 deletion.**

| gRNA | Target sequence (5'-3) | PAM |
|---|---|---|
| **gRNA1** | TCTATATGATGGTCGCTTCG | CGG |
| **gRNA2** | TGCCGAGATAAGAATTTGGG | CGG |

**Table S3. The primers used to amplify the episome, deletion target site, gRNA target site 1 and gRNA target site 2 in LER1 deletion.**

| Primers | Primers sequence (5'-3') |
|---|---|
| **Episome PF** | GTCTCTATATGATGGTCGCTTCG |
| **Episome PR** | ATACAACTTCAGAACAACGGC |
| **Primer F** | GCAGACTTACGAGACGCTATGC |
| **Primer R** | GGAAGGTCGAAGAAGGAGGC |
| **Primer F1** | GTTGACGAGGAGGGCAGTAAAC |
| **Primer R1** | CGGAGTCTGGGGAGCATG |
| **Primer F2** | TTTCAGGGTGCTAGTGCGAT |
| **Primer R2** | GGAAGGTCGAAGAAGGAGGC |

**Table S4. The primers used to identify the terminal of ΔLER1_3, ΔLER1_4, ΔLER1_7, ΔLER1_8 and ΔLER1_9.**

| Primers | Primers sequence (5'-3') |
|---|---|
| **PF** | AACCTCTTTTTAAATTGGTACCCCCTAACCCTAACCCTAACCCT |
| **Primer R3** | TATTTCTCTTCCGGTAGGGGTACCGCCCCGCTAAGCTCCTCC |
| **Primer R4** | TATTTCTCTTCCGGTAGGGGTACCGAGGGAGAGAGAGAGAAGAGGG |
| **Primer R7** | TATTTCTCTTCCGGTAGGGGTACCATGTTTGAGATAGATTCACCAGCC |
| **Primer R8** | TATTTCTCTTCCGGTAGGGGTACCATGGGAGGGATGGGATGGA |
| **Primer R9** | TATTTCTCTTCCGGTAGGGGTACCCTCGCCCTCATCTCAAGG |

**Table S5. Sequences of off-target sites identified by NGS for gRNA 1**

| | Location | Sequence (5'-3') | Off target |
|---|---|---|---|
| **Target sequence** | **20547** | TCTATATGATGGTCGCTTCGCGG | |
| **chr11** | **826542** | G.AG....TG.........A.. | N |
| **chr11** | **1139459** | .G....C.....CA...A..A.. | N |
| **chr15** | **423614** | G..T....T.C......G..A.. | N |
| **chr16** | **474644** | G.C...G....AA.......A.. | N |
| **chr1** | **415865** | ........T.TT.....GG.... | N |
| **chr24** | **55628** | ...CCT......C...C...A.. | N |
| **chr24** | **223923** | ..C...A.TA.......G..A.. | N |
| **chr25** | **259049** | ..A....T.A......CA..T.. | N |
| **chr26** | **59720** | ...G...CT.T....T....T.. | N |
| **chr4** | **1420352** | .......A..TA...GC...T.. | N |
| **chr5** | **834979** | A.G.........A..AG..G.. | N |
| **chr7** | **650804** | G..CA.A.......T.....G.. | N |
| **chr9** | **452530** | .G......G...AG....G.T.. | N |
| **chr9** | **780498** | .G...C...G..A.C.....A.. | N |

**Table S6. Sequences of off-target sites identified by NGS for gRNA 2**

| | Location | Sequence (5'-3') | Off target |
|---|---|---|---|
| **Target sequence** | **101534** | TGCCGAGATAAGAATTTGGGCGG | |
| **chr13** | **389669** | .T....A.G.T..T......G.. | N |
| **chr13** | **1036474** | .......T.C...T.G.T..... | N |
| **chr17** | **398290** | G........T...G...AA.A.. | N |
| **chr1** | **962677** | ...T...G..T.......T.... | N |
| **chr1** | **1591573** | G..G.C...........C.AG.. | N |
| **chr21** | **418030** | ...TA......A....G..CA.. | N |
| **chr22** | **104189** | .......GG.....GGA...G.. | N |
| **chr23** | **379569** | ...T..A.........GT.T... | N |
| **chr23** | **693838** | G..G....GG..C.......G.. | N |
| **chr29** | **116436** | G.A.....C..T.......AG.. | N |
| **chr2** | **552037** | .........G..GGG.A...A.. | N |
| **chr2** | **501886** | ..GG.T...TC.........A.. | N |
| **chr2** | **1192435** | .T......G..AG.......A.. | N |
| **chr2** | **1242693** | ....T..GAG..T.......G.. | N |
| **chr3** | **894992** | G...C..G.....TC.....A.. | N |
| **chr4** | **6124** | ...G.T...C....A.....A.. | N |
| **chr4** | **198614** | ...T.......A..G.G..TT.. | N |
| **chr4** | **823345** | ...AT.A.A....T......G.. | N |
| **chr4** | **832623** | .....C..C...G.....A.... | N |
| **chr7** | **684921** | ..A.....G.....A..A.AG.. | N |

| | | | |
|---|---|---|---|
| **chr7** | **617932** | ....TTC..CT.........T.. | N |
| **chr8** | **297518** | ..AG...GA..C........... | N |
| **chr9** | **214990** | ..G..T......T...GA..... | N |
| **chr9** | **1287629** | ..GG.....T...C..G...G.. | N |

**Table S7. The gRNAs for double large fragment deletions.**

| gRNA | Target sequence (5'-3) | PAM |
|---|---|---|
| **gRNA1** | TCTATATGATGGTCGCTTCG | CGG |
| **gRNA2** | TGCCGAGATAAGAATTTGGG | CGG |
| **gRNA3** | GTATGGTTGTACAATGACGG | TGG |
| **gRNA4** | TAGTGTCGTCGATTCACCAA | CGG |

**Table S8. The primers for amplification of the double large fragment deletion episome, deletion target sites, gRNA target site 1, 2, 3 and 4.**

| Primers | Primers sequence (5'-3') |
|---|---|
| **Episome PF2** | AAATGTGCCTGTTACCCTGC |
| **Episome PR2** | CGCCGTTGTTCTGAAGTTGT |
| **LER1 F** | GCAGACTTACGAGACGCTATGC |
| **LER1 R** | GGAAGGTCGAAGAAGGAGGC |
| **F1** | ATACCGCGCGGTCTTGCC |
| **R1** | CGGAGTCTGGGGAGCATG |
| **F2** | GCCCTAGATTAGTCAAGTTCGCC |
| **R2** | GAATCGGTGTTGGCTTCCAT |
| **LER2F** | CCCTATCCGTGTTGTTGCG |
| **LER2 R** | CAGCTTCTTTGCGAGGTGAAA |
| **F3** | AGTGTTTTTGCGGCATGGT |
| **R3** | GAAAAGCTTCCGCACAAATG |
| **F4** | AGGTTGCGTCACTTCACATAGA |
| **R4** | CCATCTCCATTTATCCCTCCTT |
